# Disruption of natural killer cell homing as a biomarker in persons aging with or without HIV

**DOI:** 10.1101/2022.01.05.475089

**Authors:** Kyle W. Kroll, Spandan V. Shah, Olivier A. Lucar, Thomas A. Premeaux, Cecilia M. Shikuma, Michael J. Corley, Matthew Mosher, Scott Bowler, Lishomwa C. Ndhlovu, R. Keith Reeves

## Abstract

Natural killer (NK) cells are critical modulators of HIV transmission and disease. While recent evidence suggests a loss of NK cell cytotoxicity during aging, a compound analysis of NK cell biology and aging in persons with HIV (PWH) is lacking. We set out to perform one of the first large comprehensive analyses of people aging with and without HIV to determine NK phenotypic changes during aging and how these changes are modulated while aging with HIV. Utilizing high-dimensional polychromatic flow cytometry we analyzed 30 immune-related proteins spanning broad functions such as trafficking, activation/inhibition, NK specific receptors, and memory/checkpoint receptors on peripheral NK cells from health donors, PWH with viral suppression, and viremic PWH. NK cell phenotypes are dynamic across the age span but are significantly altered in HIV and ART and with co-factors such as CMV. Specifically, NK cells in healthy aging show increasing levels of α4β7 and decreasing CCR7 expression during aging, a phenomenon nearly perfectly reversed in PWH. These HIV-associated trafficking changes could be in part due to NK cell recruitment to HIV reservoir formation in lymphoid tissue or failed mucosal signaling in the HIV-infected gut, but regardless appear to be tight biomarkers of age-related NK cell changes.

## Introduction

Advances in combination antiretroviral drug therapy (cART) and early intervention led to HIV becoming a chronic illness and an increase in life expectancy of people with HIV (PWH)(1–3). Consequently, the population of PWH are aging, with those over the age of 50 accounting for over 20% of PWH worldwide(4) and 51% in the United States(5). However, the necessity of long-term cART poses a potentially serious problem as PWH age as its impact on the immune system is still not completely understood(6). Chronic inflammation is suggested to be a leading factor of morbidity during aging(7) and PWH exhibit this chronic inflammation despite long-term viral suppression(7). Furthermore, aging PWH are at higher risk of age-related comorbidities, such as cardiovascular disease, and polypharmacy is common, which can lead to potential drug-drug interactions(8). The progression and burden of age-related comorbid conditions and multimorbidity in people aging with HIV (PAWH) differs proportionally in several ways to the general uninfected population, however the mechanisms and impact of viral directed immune responses is still not completely understood.

Natural killer (NK) cells, potent innate immune cells important in viral and tumor surveillance and immunoregulation, have been shown to play a critical role in HIV(9). Specific killer immunoglobulin-like receptors (KIRs) and human leukocyte antigen (HLA) combinations have been shown to be highly effective at control and protection from HIV infection(10, 11). Additionally, it has been shown that high NK cell functional capacity is closely associated with inhibited HIV transmission(12, 13). Interestingly, in non-human primate models of simian Immunodeficiency virus (SIV), NK cells are also shown to be highly plastic and undergo large shifts in trafficking to lymph nodes and/or gut mucosal sites(14–16). However, NK cells have also been shown to become exhausted during chronic HIV infection with typical signs being increased frequency of CD56^dim^ CD16^+^ NK cells but a decrease in functional responses(17) and a loss of Siglec-7 expression(18). Furthermore, aging has also been characterized by chronic low-grade inflammation, alterations and dysfunction in adaptive immune responses, and changes in innate immune cells(19, 20). NK cells in aging show similar dysfunctions that are seen in chronic HIV infection, namely an increase in CD56^dim^ CD16^+^ proportions but lower functional capacity(21–23) which results in an increased risk of infection(24). In addition to an increase in raw numbers of NK cells with age they also have reduced proliferative potential(21), decreased surface expression of the activating receptors NKp30 and NKp46(21), modulated KIR expression(21), loss of Siglec-7 and Siglec-9 expression(25), an accumulation of senescent cells that may be a result of age-related decline in NK cytotoxicity(26–28), and age-related trafficking changes that are directly responsible for increased susceptibility to certain pathogens(29).

To understand mechanisms underlying the interactions between NK cells, aging and HIV control, we examined NK cell phenotypic changes in PWH either on effective ART or untreated across a broad age span. We developed a high-dimensional flow cytometry comprehensive panel to measure 30 immune-parameters consisting of trafficking markers, NK cell receptors, activating/inhibitory receptors, and senescent cell markers. By defining HIV and aging specific NK cell perturbations, this will allow for the development of novel approaches to limit or reverse innate immune dysfunction, alter trajectories of co-morbidities and improve clinical outcomes in PAWH.

## Results

### Study design and cohort demographics

For our analysis we utilized a cohort of 135 samples collected by the Hawai’i Aging with HIV-1 Cohort (HAHC) at University of Hawai’i, comprised of healthy donors (HD; n = 49), PWH on treatment (cART; n = 61), and PWH off treatment (Viremic PWH; n = 25). The three groups had similar age ranges with HD having a median age of 48.01 years (32.48 – 73.48), cART having a median age of 52.99 years (26.67 – 73.34), and Viremic PWH having a median age of 42.29 years (22.28 – 78.07) (Supplemental Table 1). Sex proportion, duration of known infection, viral load, and other clinical information can be found in Supplemental Table 1. High-dimensional polychromatic flow cytometry was utilized for this study and used to quantify surface protein expression of a broad array of receptors. The panel designed covered proteins that broadly fell into (i) trafficking receptors, (ii) activation/inhibition receptors, (iii) adaptive/memory markers, and (iv) immune exhaustion markers (Supplemental Table 2).

### NK cells in aging healthy donors show paucity of receptor repertoires

To determine the impact of HIV on NK cells during aging, we first examined HIV uninfected donors (HD) in two age stratifications (under the age of 45; and over the age of 50) to establish a baseline “aged phenotype” of NK cells in the absence of known HIV infection. The median age of HD was 48.01 years old with a range of 32.48 – 73.48 years old. We focused our analyses on the dominant blood phenotype of cytotoxic CD56^dim^ CD16^+^ NK cells. Representative gating strategies for each group are shown in Supplemental Figure 1. Generalized Linear Modeling (GLM) with bootstrap resampling was employed to examine the log-odds of protein expression being a predictor of age group (Figure 1A). GLM analysis results indicated that CD127 (IL-7Rα) increase with aging is a significant predictor of aging in HD (*p* = 0.021; Figure 1A). In contrast, lower CD8α (*p* = 0.021) and CCR7 (*p* = 0.042) expression are significant predictors of younger people (Figure 1A) while KIR3DL1S1 showed a trend towards younger individuals (*p* = 0.063; Figure 1A). Interestingly, CCR7 was a significant predictor (*p* = 0.042) and negatively correlated with age (R = -0.5219, *p* = 0.0005; Supplemental Figure 2D), while the gut-homing marker α4β7 directly correlated with age (R = 0.4057, *p* = 0.0085; Supplemental Figure 2D) but was not a predictor of age in the GLM analysis (*p* = 0.978; Figure 1A). This surprising finding suggested that changes in NK cell homing repertoires may be associated with age. We also sought to utilize dimensionality reduction methods to examine the high-dimensional data on a 2-dimensional projection to interrogate whether the two age groups cluster independently. We first used Multi-Dimensional Scaling (MDS) and found that both age groups largely overlapped, highlighting that NK cells in aging remain phenotypically consistent during aging (Figure 1B). Barnes-Hut implementation of t-Distributed Stochastic Neighbor Embedding (bh-SNE) analysis further indicated that the two age groups largely overlap with minimal distinct clusters (Figure 1C). Relative expression overlaying of bh-SNE maps shows that the predictors identified previously, CD127, CD8α, and CCR7, largely do not show distinct clustering except for CCR7 which show a clearly distinct NK subset expressing high levels (Figure 1C; CD127 Top Right, CD8α Bottom Left, and CCR7 Bottom Right). We finally examined the significant predictors from the GLM analysis using violin plots (Figure 1D) to visualize expression levels between the two age groups and find that expression levels are consistent with their predictions with CD127 showing higher expression in aging while CD8α, CCR7, and KIR3DL1S1 showing decreased expression in the younger cohort. Importantly, each of these clusters and groupings supported the GLM and standard correlative analyses indicating roles for CD8α, CD127, and changes in homing for NK cells during aging.

**Figure 1.**
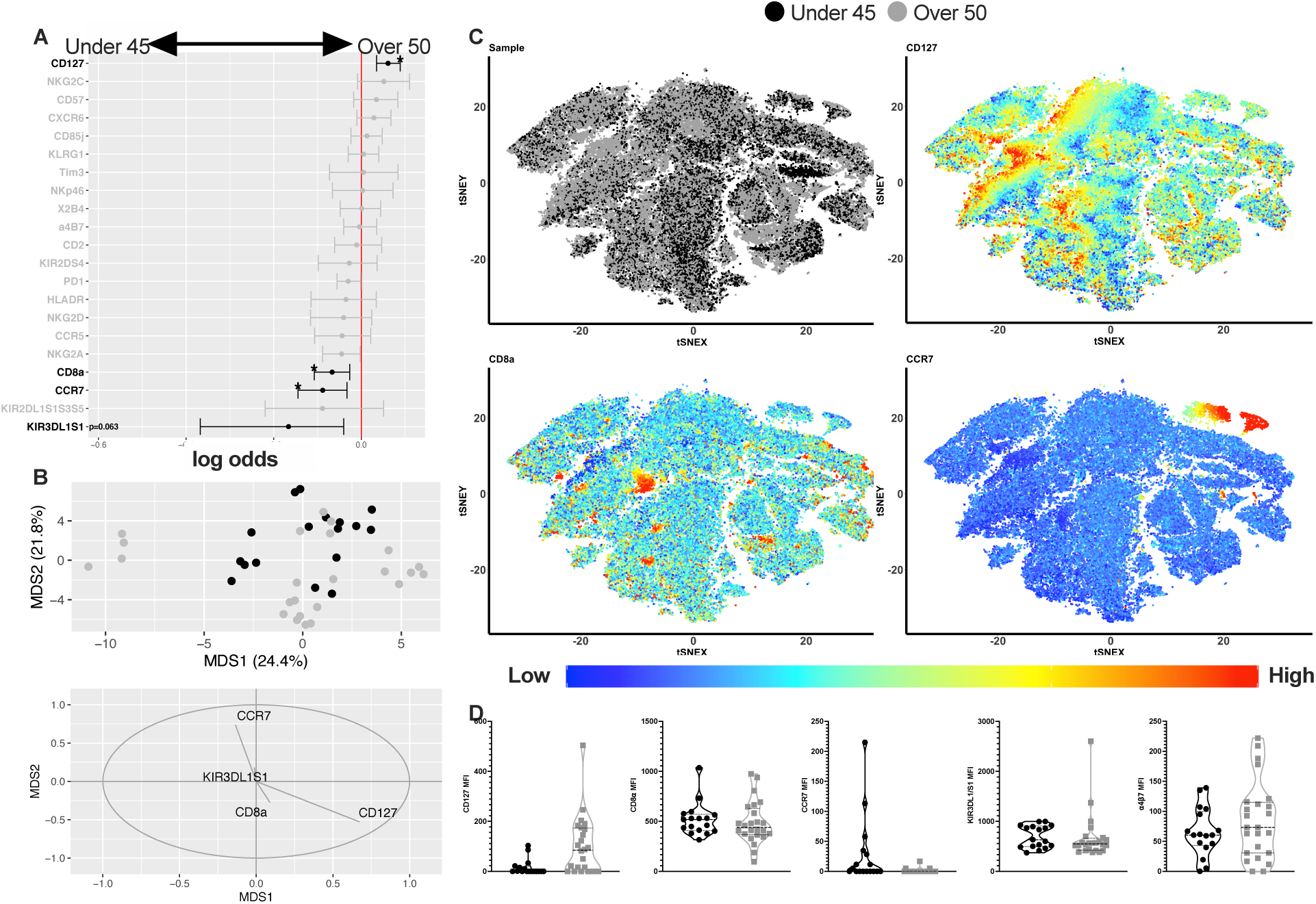
Analysis of aging on CD56^dim^ CD16^+^ NK cells. (A) Generalized linear model (GLM) with bootstrap resampling was performed to determine significant predictors of age group in HD. (B) Multidimensional scaling (MDS) was performed to determine clustering in dimensionality reduced projection. Black circles represent HD under the age of 45 and grey circle represent HD over the age of 50. MDS loadings are displayed for the significant predictors from A. (C) t-Distributed Stochastic Neighbor Embedding (t-SNE) was performed using sampling approach from A. Relative marker expression overlays use colorimetric scale from low (blue) to high (red) relative expression of the protein. (D) Violin plots of median fluorescence intensity (MFI) for significant predictors of age group. *BH adjusted *p* < 0.05

### NK cell aging phenotypes in PWH diverge in people over 50 compared to uninfected controls

Next, we sought to determine the impact of HIV infection on the NK cell aging phenotype seen in HD. We examined two groups of PWH: those that were either on cART or not on treatment (Viremic) at the time of sampling (Supplemental Table 1). Using the same GLM with bootstrap resampling as before, we first examined PWH compared to HD for those under the age of 45. Surprisingly, there were few significant predictors of HIV status in this age group (Figure 2), with only PD-1 and HLA-DR being significant when HD were compared to cART (p *p* = 0.021 and *p* = 0.021, respectively; Figure 2A). When comparing HD and Viremic under the age of 45, NKG2C and α4β7 were potential predictors of HIV (*p* = 0.084 and *p* = 0.084, respectively; Figure 2B) status while PD-1 was a predictor of the HD group (*p* = 0.042; Figure 2B). Interestingly, α4β7 had a significant negative correlation with age in both the cART (Supplemental Figure 3D) and Viremic groups (Supplemental Figure 4D), which is inverse to the correlation seen in HD aging (Supplemental Figure 2D), again suggesting that aging with HIV causes a shift in NK cell trafficking. In contrast to the under 45 group, many proteins emerged as significant predictors of both HIV groups for the over 50 group. For PWH on cART over the age of 50, CD2, CCR7, and α4β7 emerged as significant predictors compared to HD (*p* = 0.014, *p* = 0.014, and *p* = 0.031, respectively; Figure 2C). These predictors were consistent in the Viremic group (*p* = 0.021, *p* = 0.021, *p* = 0.021, respectively; Figure 2D), in addition the proteins CCR5, CD85j, CCR7, and HLA-DR were also potential predictors of Viremic PWH compared to HD (*p* = 0.0945, *p* = 0.021, *p* = 0.021, and *p* = 0.084, respectively; Figure 2D). All together these results again strongly indicate NK cell trafficking is modulated in HIV and aging, as shown by a decreased expression of the gut-homing marker α4β7 and increased expression of the lymph node homing marker CCR7. Broadly this change could be considered a reversal of the expected healthy aging homing repertoire of NK cells.

**Figure 2.**
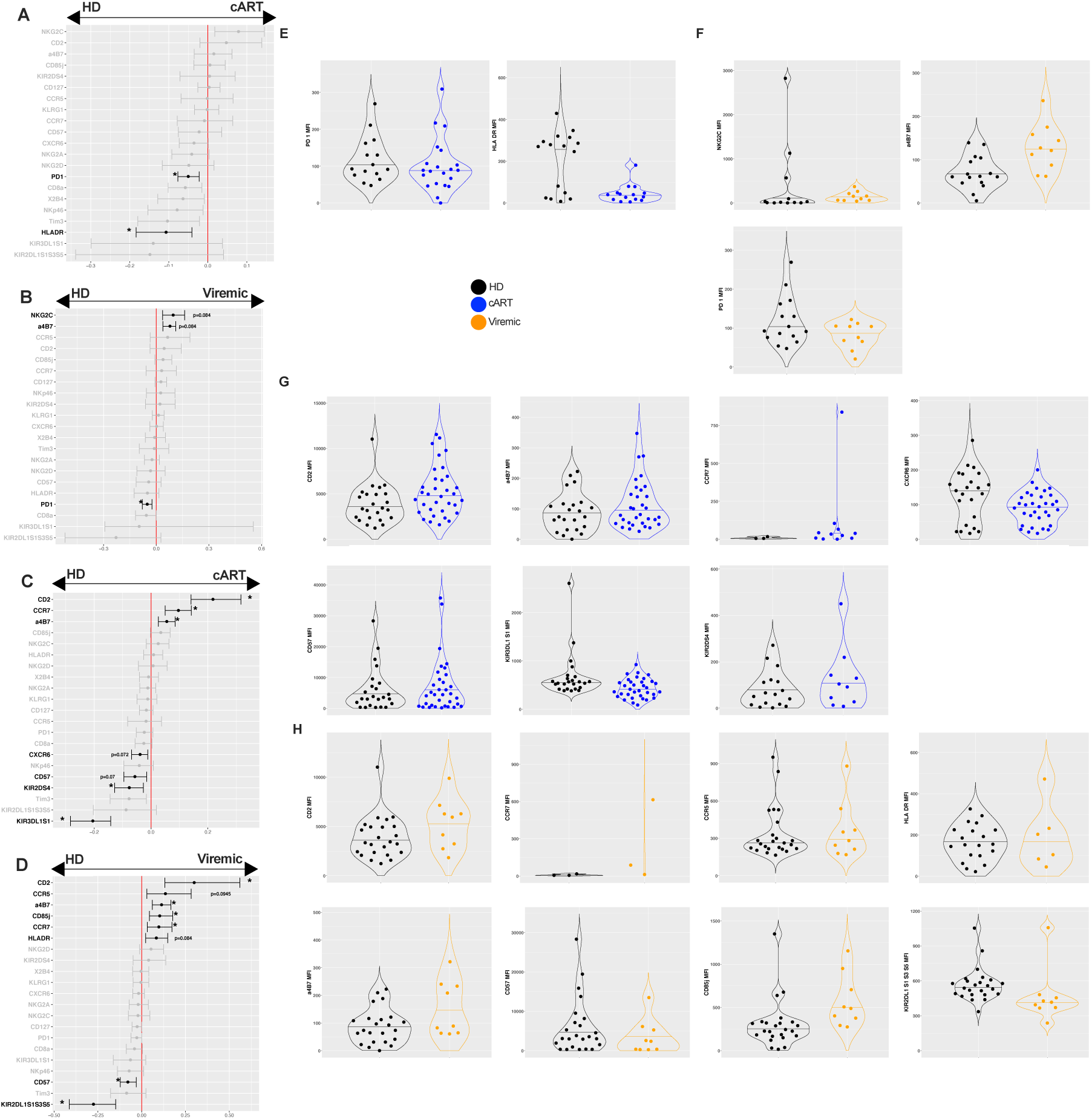
GLM with bootstrap resampling was used to determine predictors of (A) HD and cART under the age of 45; (B) HD and Viremic PWH under the age of 45; (C) HD and cART over the age of 50; and (D) HD and Viremic PWH over the age of 50. Violin plots show expression levels of highlighted markers. (E) HD and cART under 45. (F) HD and Viremic PWH under 45. (G) HD and cART over 50. (H) HD and Viremic PWH over 50. Black circles represent HD, blue circles represent cART, and orange circles represent Viremic PWH. *BH adjusted *p* < 0.05

### HIV viral load and duration of known infection predicts NK cell repertoires in persons aging with HIV

NK cell changes in functional capacity, phenotype, exhaustion, and trafficking have been previously shown to correlate with viral load(30–34). Due to these previously described correlations, we next aimed to evaluate potential correlates among HIV viral load and observed NK phenotypes in the Viremic group, independent of age. Interestingly, viral load did not correlate with NK cell frequency (Figure 3A), activation/checkpoint markers (Figure 3B), nor NK cell receptors (Figure 3C). Surprisingly, only CCR7 significantly correlated with viral load (R = 0.4804, *p* = 0.0275; Figure 3D). None of the other trafficking markers showed significant correlations; however, α4β7 did show a positive trend with viral load (R = 0.3758, *p =* 0.0932; Figure 3D). These analyses reenforce the fact that even independent of aging, HIV infection and ongoing virus replication influences NK cell homing and potential tissue localization.

**Figure 3.**
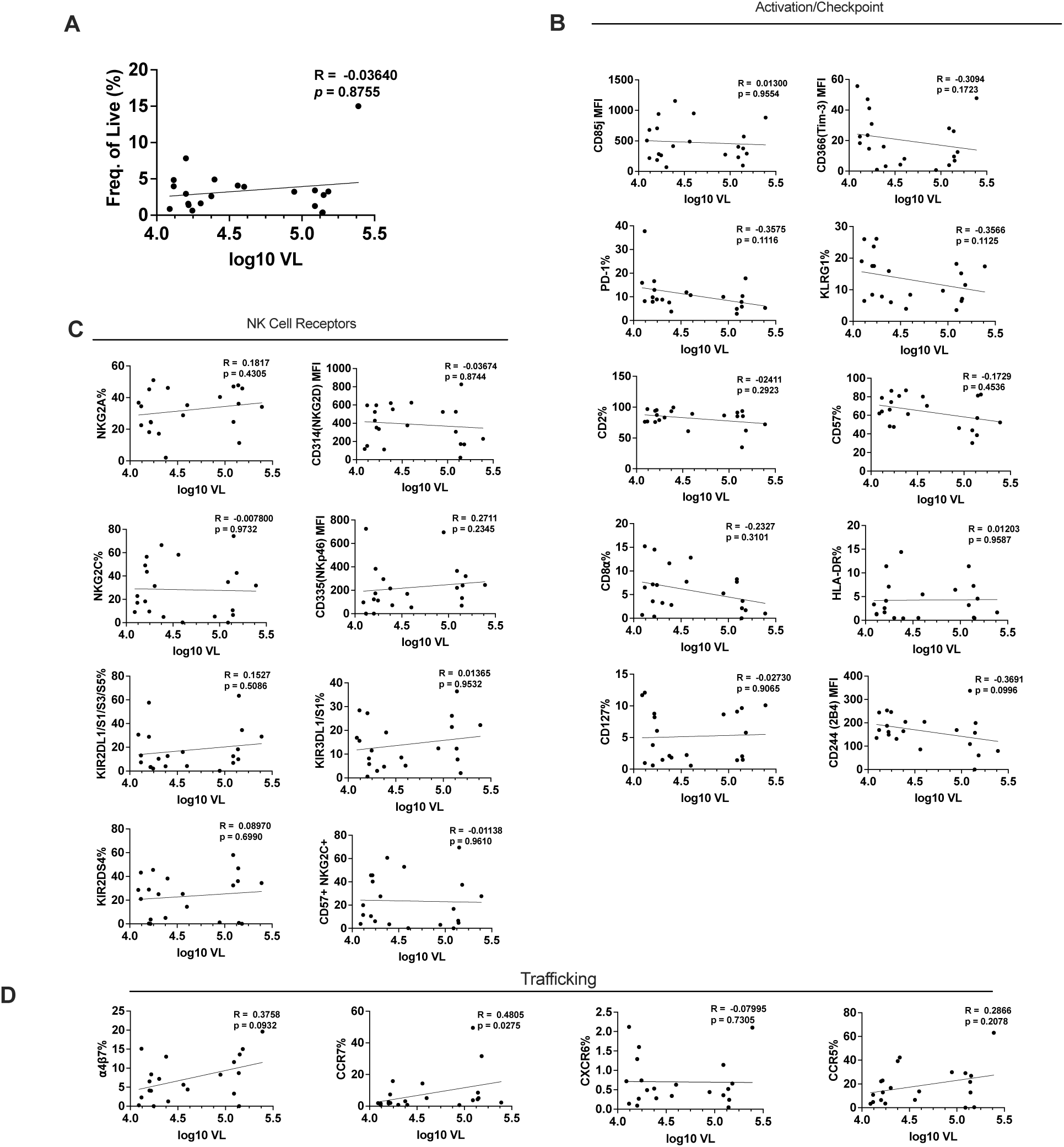
Correlation analysis of HIV viral load and protein expression levels on CD56^dim^ CD16^+^ NK cells in PWH. (A) CD56^dim^ CD16^+^ NK cell frequency of live lymphocytes. (B) Activation and/or checkpoint receptors. (C) NK cell receptors including NCRs and NKG2 family receptors. (D) Trafficking receptors including mucosae-homing marker α4β7 and lymph-node homing marker CCR7. Spearman R and *p*-values noted on each graph.

Given the duration of HIV infection may play a role in altered NK cell phenotype in PWH, we performed correlation analysis between NK cell markers and duration of known infection for PWH with or without cART (Figure 4). Unexpectedly, a significant positive correlation with duration of known infection and frequency of NK cells in PWH on cART was observed (Figure 4A). Interestingly, only 2B4 showed a positive trend with duration of infection among Viremic PWH (R = 0.3918, *p* = 0.0790; Figure 4B). NK specific receptors did not correlate with duration of infection, except for NKp46 in the cART group showing a weak negative correlation (Figure 4C), and correlations between duration of known infection and NK trafficking were not observed (Figure 4D). Collectively, these data suggest duration of infection could influence the NK cell repertoire, but rather age and viremia are predictors of the HIV-associated trafficking change.

**Figure 4.**
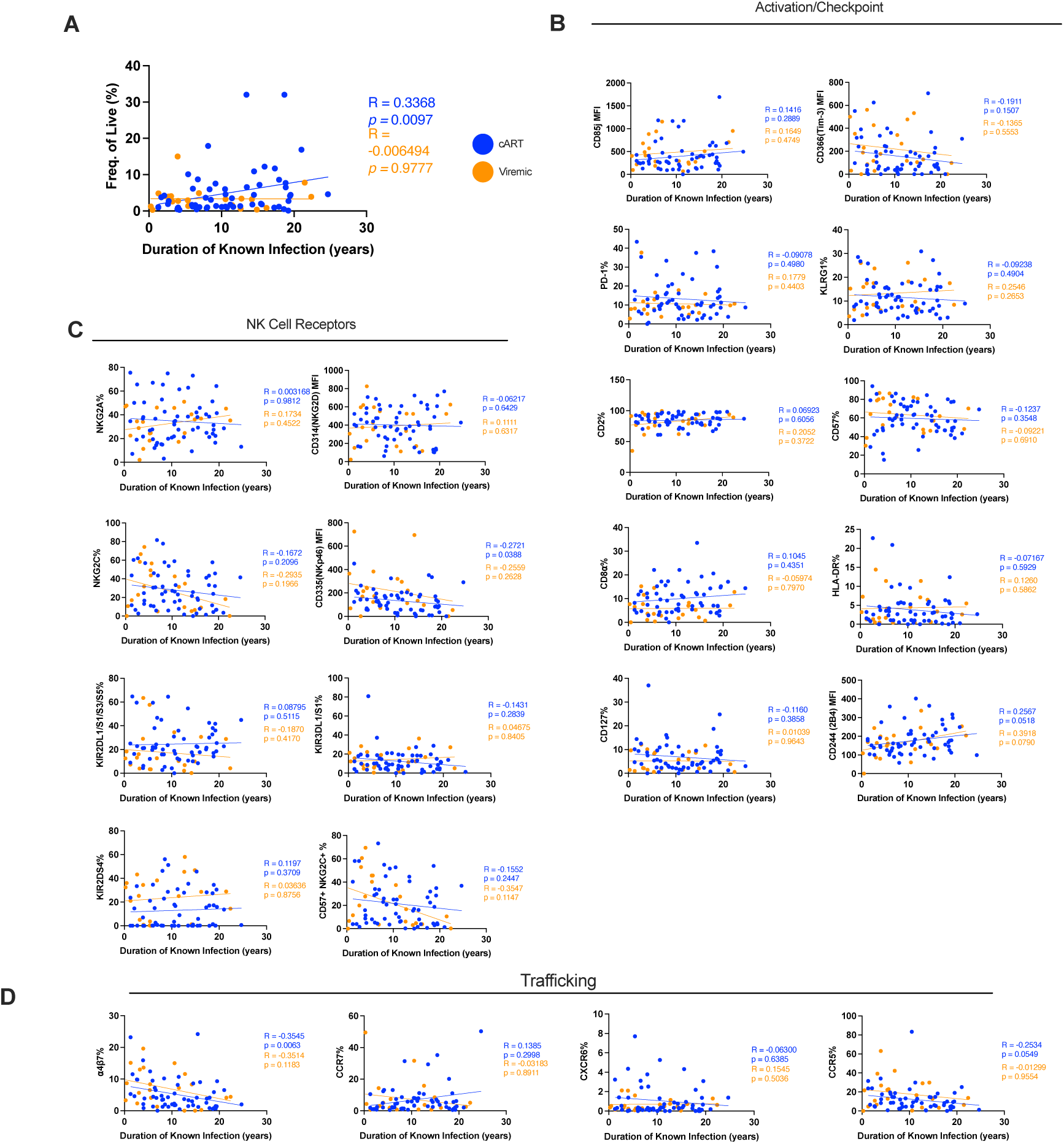
Correlation analysis of duration of known infection in PWH and NK protein expression. (A) CD56^dim^ CD16^+^ NK cell frequency of live lymphocytes. (B) Activation and/or checkpoint receptors. (C) NK cell receptors including NCRs and NKG2 family receptors. (D) Trafficking receptors including mucosae-homing marker α4β7 and lymph-node homing marker CCR7. Blue circles, lines, and text represent results of cART. Orange circles, lines, and text represent results of Viremic PWH. Spearman correlation was used for determining correlation and significance.

### Elevated CMV IgG titers is linked to higher expression of several NK cell aging phenotypes in those over 50

NK cells are known to undergo phenotypic and functional changes following CMV infection, mainly through the expansion of an adaptive NK pool that are CD57^+^ NKG2C^+^ (35–37). To determine if age-related changes observed are due to CMV infection we evaluated plasma anti-CMV IgG antibody titers in all cohort groups. Consistent with known data on CMV and HIV co-infections(38), CMV antibody titers were elevated in PWH compared to HD (Supplemental Table 1). The CMV+ HD over 50 years of age had higher proportions of NK cells expressing CD57, NKG2C, CD127, PD-1, and CD85j (Figure 5A), which could represent an expanded adaptive NK subset previously described by our group and others(35-37,39). Interestingly, we did not see a significant impact of CMV on α4β7 or CCR7 in HD (Figure 5B-5D). However, increased CCR5 expression is observed with CMV titers in both HD age groups (Figure 5E) and positively correlates with age (Supplemental Figure 5D). Overall, these data suggest that CMV is a modulator of adaptive NK cells and other NK cell receptors, but changes in PWH are largely dominated by age and HIV status (Supplemental Figure 8). Specifically, HIV and age drive the CCR7/α4β7 trafficking change independent of CMV (Supplemental Figures 3,4 6-8). Even with the limited data within range we do see a significant negative correlation in the cART group between CXCR6 expression and CMV IgG titer (R = -0.3721, *p* = 0.0072; Supplemental Figure 6D). However, we do not see the same correlation in the Viremic group for CXCR6 (R = 0.3404, *p* = 0.1311; Supplemental Figure 7D). Chord plots are consistent with these findings as well, showing that it appears that HIV is the main driver of NK phenotypic changes in both the cART and Viremic groups (Supplemental Figure 8).

**Figure 5.**
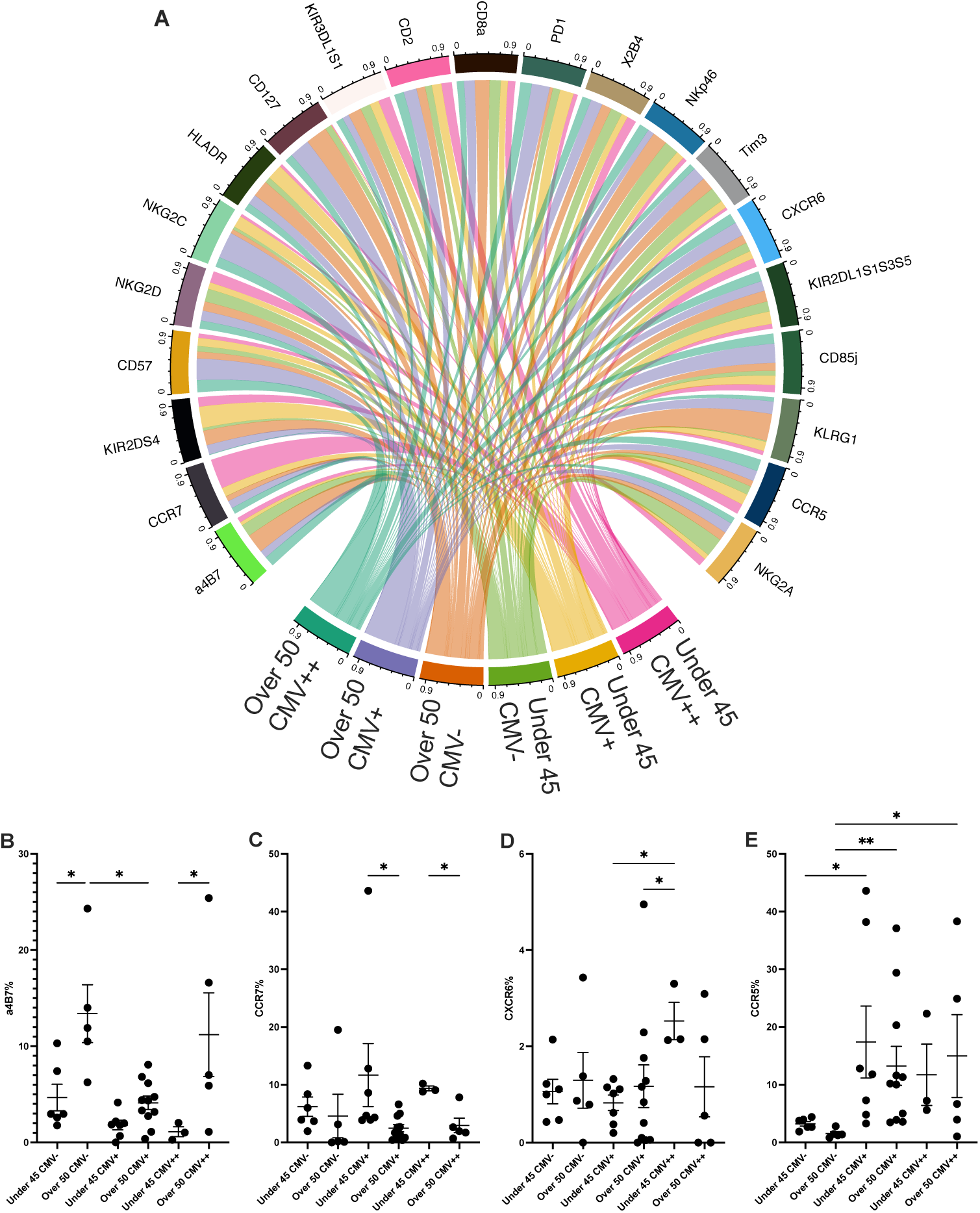
Analysis of NK protein expression based on age and CMV status. (A) Chord plot showing proportion of median fluorescence intensity for all markers in HD, delineated by age group and CMV titer. Expression levels of trafficking markers in HD based on CMV titer comparing the expression of α4β7 (B), CCR7 (C), CXCR6 (D), and CCR5 (E). * p < 0.05; ** p < 0.01. Kruskal-Wallis test followed by Benjamini Hochberg FDR correction for multiple testing.

## Discussion

Delineating meaningful changes in NK cell profiles during aging and in PWH has remained poorly understood, limiting a complete understanding of the cellular changes provoked following infection that may result in subsequent secondary disease states. To help minimize this gap we used 28-color polychromatic cytometry to robustly characterize cell surface changes in the NK cell repertoire of healthy persons and persons aging with HIV on and off cART. Indeed, we found one of the tightest predictors of how the global NK cell repertoire changes during healthy aging can be predicted solely by increasing α4β7 expression. Furthermore, we find a total reversal of this phenomenon in PWH.

Aging has been found to increase the CD56^dim^ CD16^+^ of NK cells while decreasing functional capacity and even further contributing to systemic inflammation(24). Our data demonstrated a well conserved aging NK phenotypic profile consisting of NK cell specific, activation and altered checkpoint specific receptors. Interestingly, a highly distinctive trafficking pattern was identified with NK cells in aging consisting of decreased lymph node homing via downregulation of CCR7, in conjunction with increased gut mucosae homing via upregulated α4β7. Previous reports have identified age-associated dysfunction in the gut integrity resulting in microbial translocation and macrophage dysfunction(40). Our data could suggest that increased recruitment of NK cells to the gut mucosae could be one mechanism for maintaining gut homeostasis during aging. Further investigation into the cytolytic capabilities of trafficked NK cells to the gut mucosae can provide more robust insights into the identified trafficking repertoire.

Introduction of cART therapy has dramatically lengthened the lifespan of PWH, resulting in a novel cohort crucial to understanding HIV pathogenesis. One of the most interesting findings in this study was the modulation of NK cell trafficking pattern with HIV infection. Regardless of cART treatment, NK cells from PWH showed a significant decrease in *α*4*β*7 expression with a trend towards increased CCR7, potentially indicating a reduction of NK cell trafficking to the gut, but instead mobilizing to the lymph nodes, perhaps due to lymphoid follicles being reservoirs for both HIV and SIV(41). Despite the effectiveness of cART, HIV can survive in follicular helper T cells(42–44), remaining undetected by immunoregulatory activation. Using non-human primate models, previous reports have compared the follicular regions in chronically infected SIV pathogenic and non-pathogenic hosts and saw an increase of NK cell recruitment in elite controllers(15, 45). It is possible that in PWH, NK cells are mobilizing to the lymph nodes in attempt to clear residual HIV infected T cells.

Another component of our study was to compare putative biomarkers for aging NK cells in infected and uninfected individuals under 45 or over 50. Surprisingly, we found that younger persons had NK cell repertoires that are relatively similar regardless of HIV status, but noticeable disparities became clearly apparent in those over the age of 50. CD2, CCR7, and *α*4*β*7 were significant predictors compared to healthy donors for PWH on cART, with the addition of CCR5 and CD85j for untreated PWH. Our data suggests that neither aging nor HIV alone severely impacts the NK cell repertoire, but perhaps rather a combination of the two does. Aging and HIV are associated with a variety of similar cellular perturbations relating to function, proliferation, and exhaustion(23). A comparison of elderly HIV-infected individuals to healthy donors in the same age cohort found increased rates of hypertension, hypertriglyceridemia, and other disorders in HIV-infected individuals(46). It is possible that a combination of the two have a synergistic and significant effect on the NK cell phenotypic profile. Altogether, these data highlight the age-associated NK cell phenotypic changes in HIV-infected adults and their potential clinical implications.

In line with previous research, we identified NK cells from CMV infected aged adults to have higher expression of CD57 and NKG2C(35,36,47), a unique subset of adaptive NK cells primed by the virus. Although we found age-related NK cell phenotypic changes also associated with CMV status, there was minimal impact of CMV infection specifically on NK trafficking receptor expression in healthy donors. Specifically, there was no significant difference between CMV+ and CMV- aged individuals for NK cell *α*4*β*7, CCR7, or CXCR6. These data suggest that age-related changes in the NK cell repertoire do occur due to CMV infection but are largely independent from those induced by HIV.

Importantly, we acknowledge several limitations in our study which were predominantly focused on limited treatment information and sample quality issues. Due to the age or other limitations of some samples, we were unable to obtain CMV viral shedding from cryopreserved urine samples. In addition, treatment history, specific antiretroviral drugs used, and any potential interruptions in treatment were not known for all participants. Additionally, as many study participants were on ART regimens that are no longer common care, the impact of other drugs on NK cells and chronic inflammation will need to be further examined.

Altogether, we provide an aging NK cell phenotypic profile, delineate the modulations induced by HIV infection, and highlight the combined dysfunctional properties of HIV and aging. To our knowledge, this is the first study of NK cell phenotypes in aging PWH in both treated and untreated aged cohorts. NK cells have been linked to the control and disease progression of HIV making it imperative to understand the natural trafficking patterns of these cells and the subsequent modulations following HIV infection. Further research will be needed to evaluate the functional capabilities of aged NK cells in PWH on and off cART and the consequences of these changes.

## Materials and Methods

### Flow cytometry

Flow cytometry staining of PBMCs was performed using a 28-color surface phenotype panel (Supplemental Table 2). Briefly, cryopreserved PBMCs were rapidly thawed at 37 °C and immediately transferred to complete media pre-warmed to 37 °C. After centrifugation, PBMCs were incubated for 30 minutes at 4 °C in Blue Live/Dead (Invitrogen, Carlsbad, CA). PBMCs were then washed and surface stained for 20 minutes at room temperature. Samples were then stained immediately after with Streptavidin- BUV395 as a secondary antibody for biotinylated CD159c for 15 minutes at RT. PBMCs were washed once more and fixed with 2% paraformaldehyde. All acquisitions were carried out on a FACS Symphony cytometer (BD Biosciences) and analyzed with FlowJo v10.7.1 (BD Biosciences). The gating strategy used to define NK cells was as follows: Live, Lineage^-^ (CD19^-^ CD14^-^ CD4^-^ CD3^-^) CD56^dim^ CD16^+^.

### CMV antibody titer quantification

Matched cryopreserved plasma samples were used for quantification of Human Cytomegalovirus IgG and IgM antibody titers (Quest Diagnostics; order code 6732; Marlborough, MA, USA).

### HIV viral load quantification

Plasma viral loads were assessed using Amplicor HIV-1 Monitor Ultrasensitive Assay (Roche Molecular System, Branchburg, NJ).

### Multidimensional analyses

Multidimensional scaling (MDS)(48) was performed using the CytoGLMM R package as described by package documentation(49) using median summarized expression values for each sample. Uniform Manifold Approximation and Projection (UMAP)(50) was performed with a beta version of the CytoDRAV(51) application that implements the uwot v0.1.10 R package(52). Input data were transformed using the inverse hyperbolic sine function with a cofactor of 5. Default parameters were used and results visualized with the ggplot2 R package(53).

### Statistical analysis

Spearman correlation analysis was performed in Prism v9.0 (GraphPad Software). Generalized linear modeling (GLM) with bootstrapping was performed with the CytoGLMM R package(54) with R v3.6.3(55). GLM with bootstrapping was performed according to previously published studies(56, 57). Briefly, compensated FCS files of CD56^dim^ CD16^+^ NK cells were exported from FlowJo and loaded into R. Data were transformed using the hyperbolic sine transformation with a cofactor equal to 5 and a random sampling of 2000 cells from each sample, or all cells if the sample contained fewer than 2000 cells, was performed. The *cytoglm* function of the CytoGLMM package was used to perform 1000 iterations of bootstrap resampling with replacement followed by logistic regression. Results are reported as log-odds of a given marker predicting binary group assignment (HD and ART; HD and PWH; HD under 45 and HD over 50). Corrections for multiple testing were performed using the Benjamini-Hochberg method for controlling false discovery rate(58). P-values reported are BH adjusted p-values. An adjusted p-value cutoff of 0.05 was used to determine significance. Median fluorescence intensities (MFI) were exported from FlowJo for comparing marker expression between groups. These data were not transformed prior to statistical analysis. Mann-Whitney *U*- tests were used to compare MFI levels between groups.

### Study approval

Cryopreserved human peripheral blood mononuclear cells (PBMCs) were obtained from the Hawai’i Aging with HIV-1 Cohort (HAHC) at University of Hawaii. Details of the HAHC study enrollment and clinical characterization were previously published(59) and approved by the University of Hawai’i Institutional Review Board. All participants signed institutional review board–approved informed consent forms prior to participation. CD4 lymphocyte counts were obtained in real-time by standard technique from a local CAP certified reference laboratory. Demographic and clinical details of study participants are provided in Supplemental Table 1.

## Acknowledgements

This research was supported by National Institutes of Health (NIH) grants: R01AI120828, R01AI143457, R01AI161010 (to R.K.R.). We also acknowledge support from the CVVR Flow Cytometry and Harvard University Center for AIDS Research Advanced Laboratory Technologies Core (P30 AI060354). The authors thank Michelle Lifton for technical assistance and panel design in flow cytometry.

## Author Contributions

L.C.N. and R.K.R. designed the study. K.W.K., S.V.S., O.A.L., T.A.P., M.J.C., M.M., and S.B. performed the experiments and analyzed data. K.W.K. performed the bioinformatic analyses. C.M.S. provided samples critical to the study. All authors contributed to the writing of the manuscript.

