## Supplemental Material for "Disruption of natural killer cell homing as a biomarker in persons aging with or without HIV"

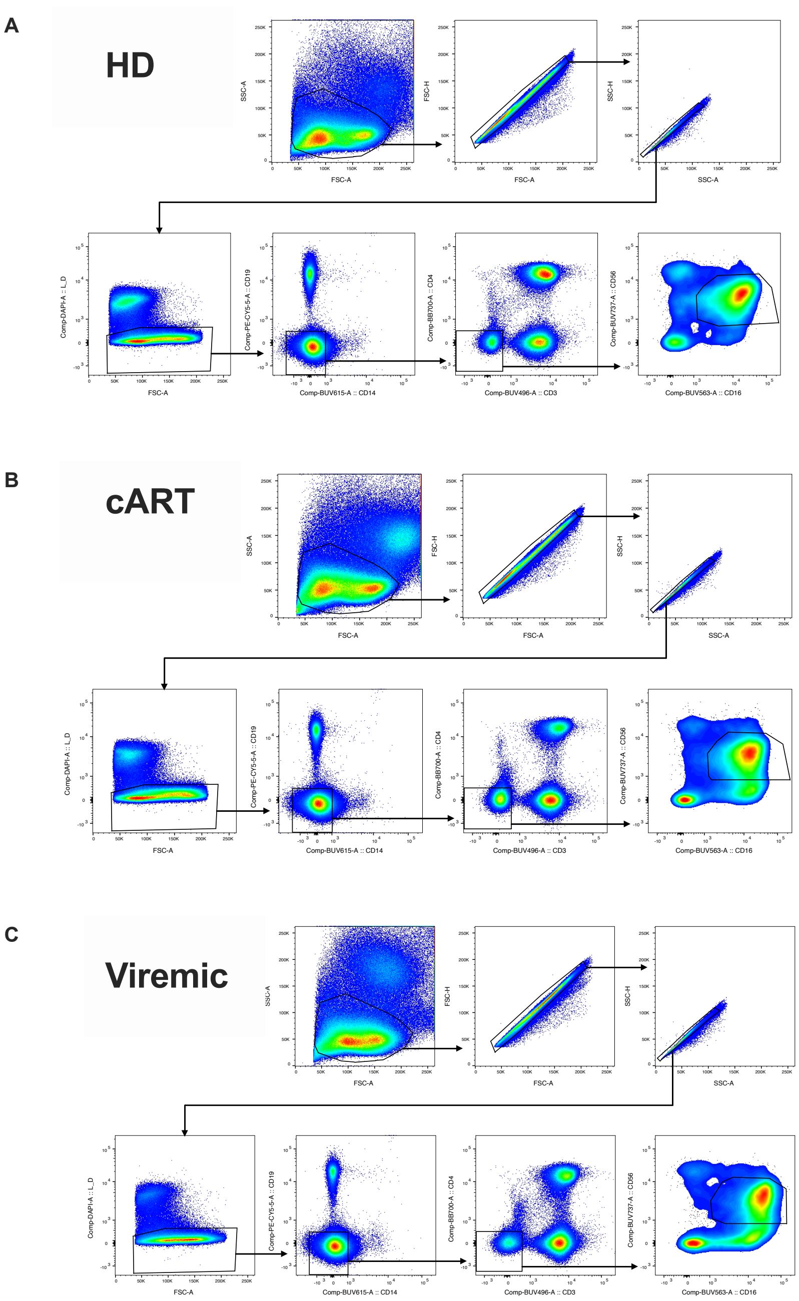


**Supplemental Figure 1.** Gating strategies used to define CD56dim CD16+ NK cells in (A) HD, (B) cART, and (C) Viremic.

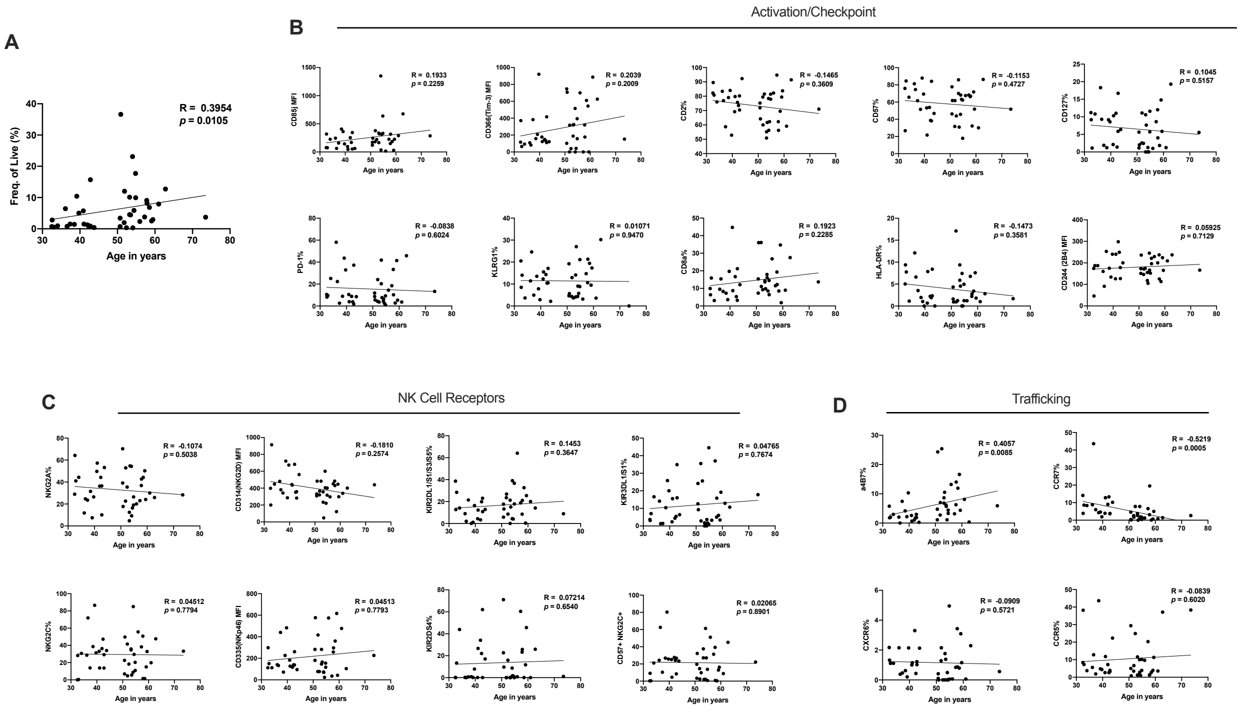


**Supplemental Figure 2.** Correlations of NK cell receptors and age in HD. Correlations between age and (A) frequency of CD56$^{dim}$ CD16$^{+}$ NK cells, (B) activation/checkpoint receptors, (C) NK cell specific receptors, and (D) trafficking receptors. Spearman correlation values and p-values are reported within each subfigure.

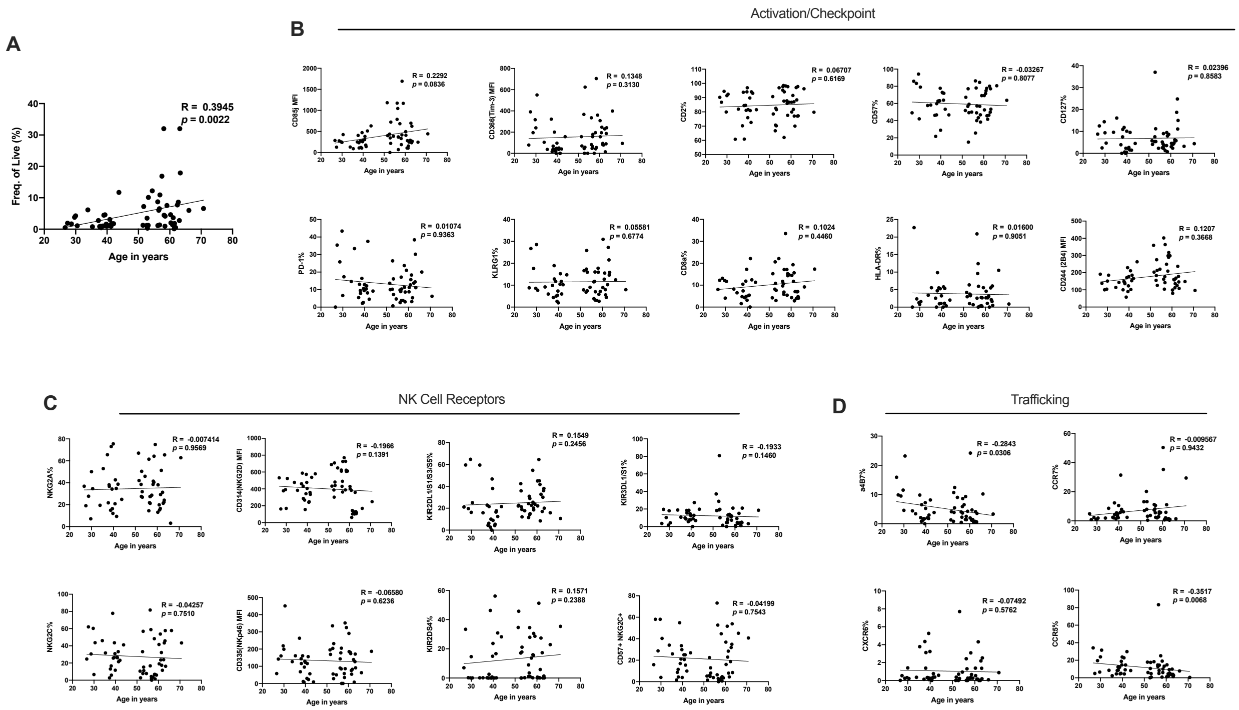


**Supplemental Figure 3.** Correlations of NK cell receptors and age in cART. Correlations between age and (A) frequency of CD56$^{dim}$ CD16$^{+}$ NK cells, (B) activation/checkpoint receptors, (C) NK cell specific receptors, and (D) trafficking receptors. Spearman correlation values and p-values are reported within each subfigure.

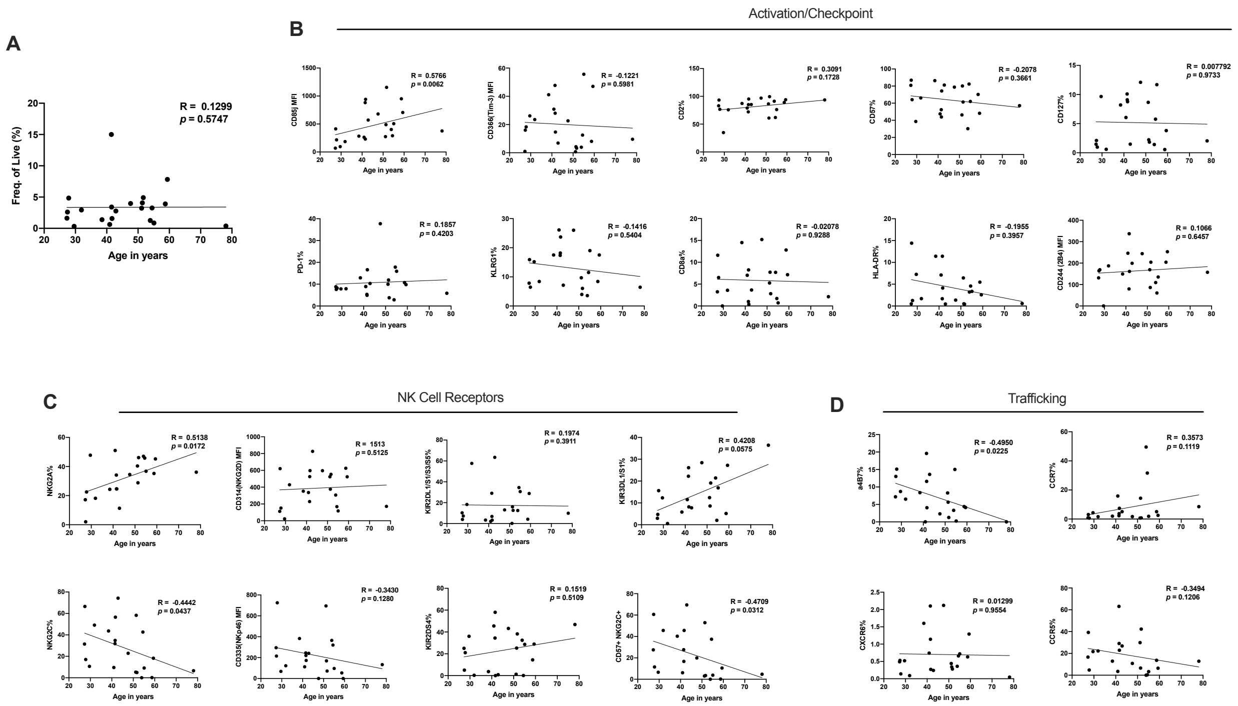


**Supplemental Figure 4.** Correlations of NK cell receptors and age in Viremic PWH. Correlations between age and (A) frequency of CD56$^{dim}$ CD16$^{+}$ NK cells, (B) activation/checkpoint receptors, (C) NK cell specific receptors, and (D) trafficking receptors. Spearman correlation values and p-values are reported within each subfigure.

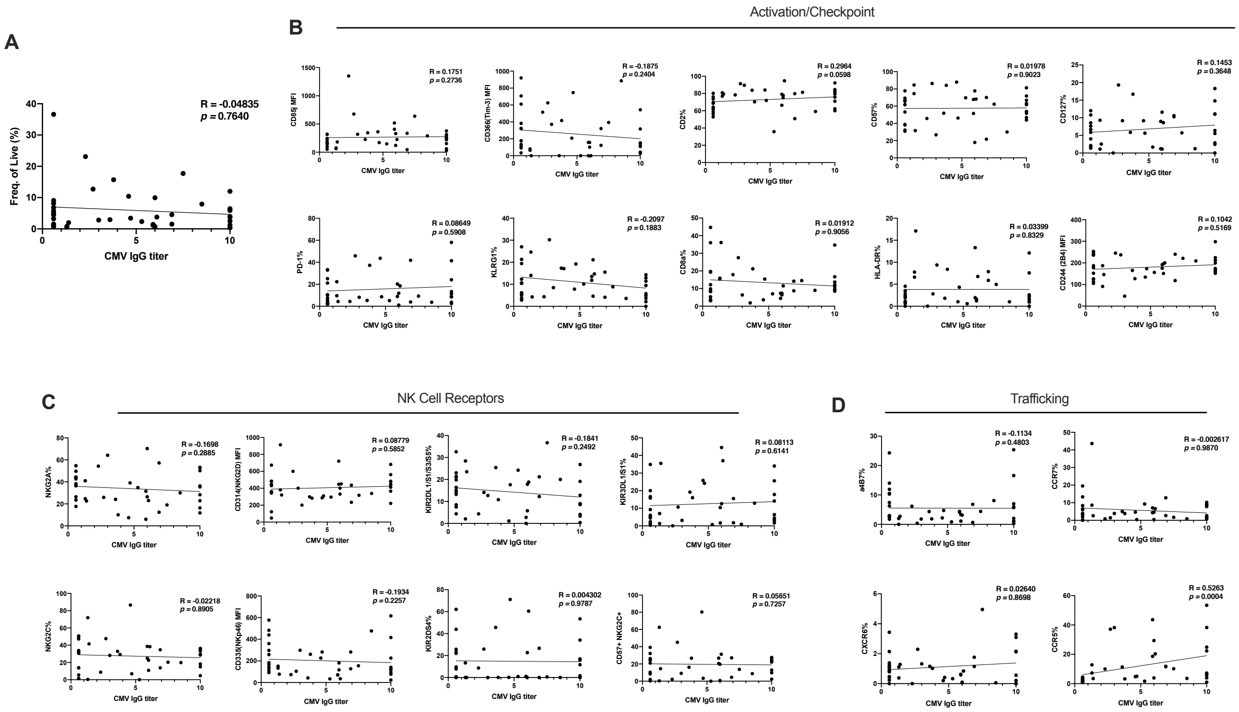


**Supplemental Figure 5.** Correlations of NK cell receptors and CMV IgG titers in HD. Correlations between CMV IgG titers and (A) frequency of CD56$^{dim}$ CD16$^{+}$ NK cells, (B) activation/checkpoint receptors, (C) NK cell specific receptors, and (D) trafficking receptors. Spearman correlation values and p-values are reported within each subfigure.

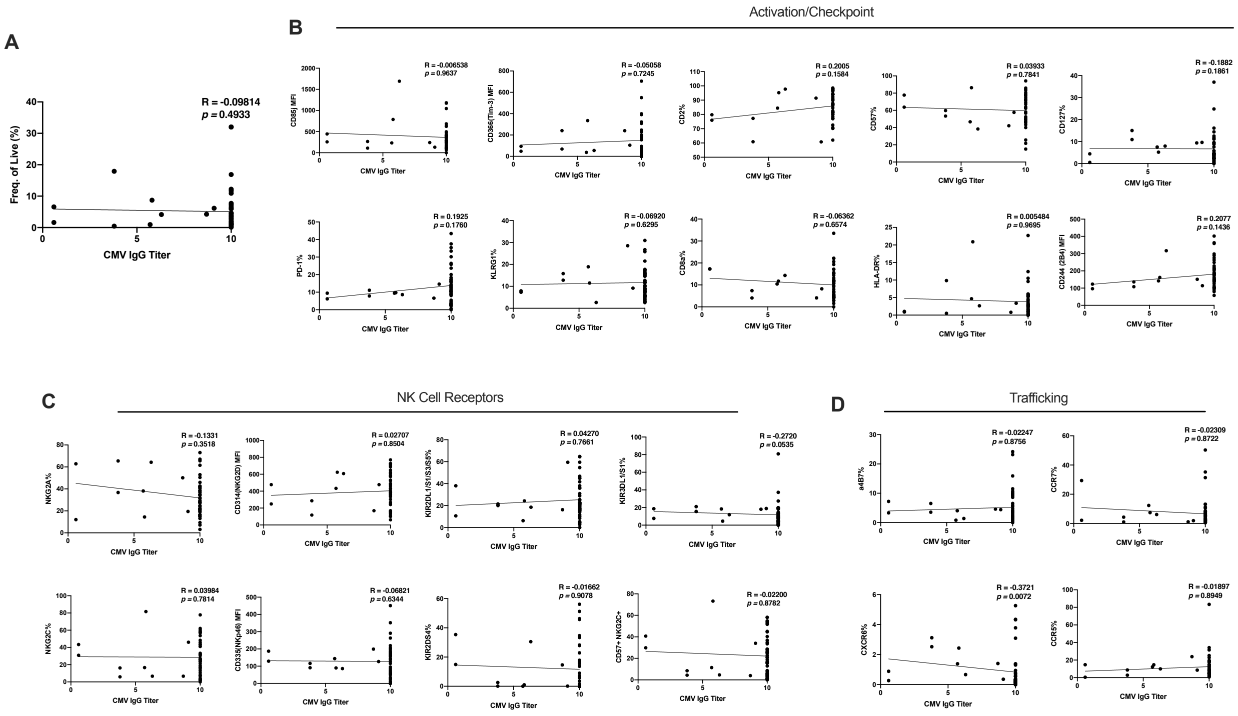


**Supplemental Figure 6.** Correlations of NK cell receptors and CMV IgG titers in cART. Correlations between CMV IgG titers and (A) frequency of $CD56^{dim}$ $CD16^{+}$ NK cells, (B) activation/checkpoint receptors, (C) NK cell specific receptors, and (D) trafficking receptors. Spearman correlation values and p-values are reported within each subfigure.

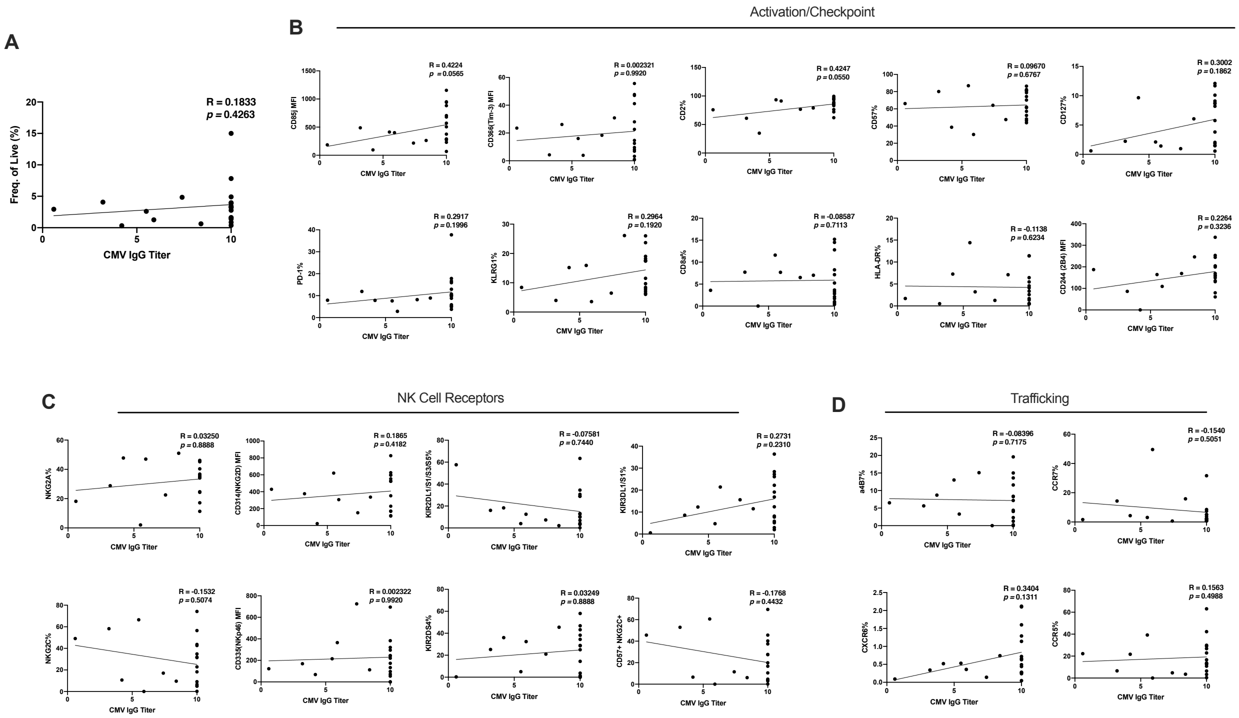


**Supplemental Figure 7.** Correlations of NK cell receptors and CMV IgG titers in Viremic PWH. Correlations between CMV IgG titers and (A) frequency of $CD56^{dim}$ $CD16^{+}$ NK cells, (B) activation/checkpoint receptors, (C) NK cell specific receptors, and (D) trafficking receptors. Spearman correlation values and p-values are reported within each subfigure.

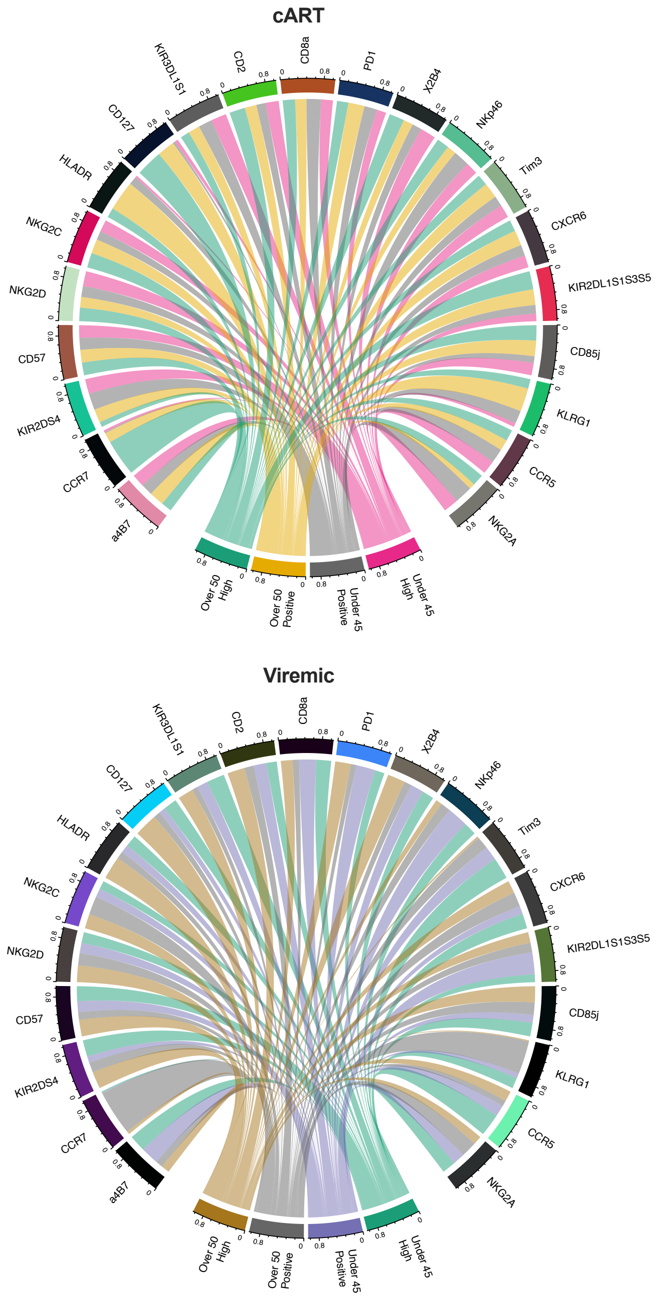


**Supplemental Figure 8.** Chord plots of proportion of MFI expression for receptors in both cART and Viremic PWH. Bands represent proportion of MFI that is explained by connecting CMV group.

| | HD | PWH on cART | PWH |
|---|---|---|---|
| Age, years: Median (Range) | 48.01 (32.48-73.48) | 52.99 (26.67-73.34) | 42.29 (22.28-78.07) |
| Sex: proportion male | 39/49 | 52/62 | 18/24 |
| Duration of known infection, years: Median (range) | N/A | 10.55 (1.29-24.67) | 6.24 (0.25-22.38) |
| Viral load, copies/ml: Median (range) | N/A | <50 | 30551.5 (12300-245000) |
| CD4: count/ml Median (range) | N/A | 528 (132-1380) | 332 (4-751) |
| NRTI total exposure, years: Median (range) | N/A | 6.48 (0.08-19.12) | 2.25 (0-13.42) |
| NNRTI total exposure, years: Median (range) | N/A | 1.88 (0-9.63) | 0 (0-6.25) |
| PI total exposure, years: Median (range) | N/A | 1.99 (0-9.61) | 0.12 (0-5.50) |
| Cytomegalovirus IgG Antibody Titer: Median (range) | 3.80 (<0.60 - >10.00) | >10.00 (< 0.60 - >10.00) | >10.00 (< 0.60 - >10.00) |

**Supplemental Table 1.** Cohort demographic and clinical information

| Target | Fluorochrome | Clone | Manufacturer |
|---|---|---|---|
| LIVE cells | Blue Live/Dead | | Invitrogen |
| CD19 | PE-CY5.5 | J3-119 | BECKMAN COULTER |
| CD14 | BUV615 | M5E2 | BD Pharmingen |
| CD3 | BUV496 | UCHT1 | BD Pharmingen |
| CD4 | BB700 | L200 | BD Pharmingen |
| CD56 | BUV737 | NCAM16.2 | BD Pharmingen |
| CD16 | BUV563 | 3G8 | BD Pharmingen |
| NKP46 | BV711 | 9E2/NKp46 | BD Pharmingen |
| NKG2D | BB790 | 1D11 | BD Pharmingen |
| NKG2C | Biotin + SA BUV395 | REA205 | MILTENYI |
| NKG2A | PE-CY7 | Z199 | BECKMAN COULTER |
| KIR3DL1/S1 | VioBlue | REA168 | MILTENYI |
| KIR2DL1/S1/S3/S5 | FITC | HP-MA4 | BIOLEGEND |
| KIR2DS4 | APC-Vio770 | REA284 | MILTENYI |
| CD85J | PE | GHI/75 | BD Pharmingen |
| a4b7 | APC | | NHP Reagent Resource |
| CXCR6 | BV786 | 13B 1E5 | BD Pharmingen |
| CCR5 | PE-CY5 | 2D7/CCR5 | BD Pharmingen |
| CCR7 | APC-R700 | 3D12 | BD Pharmingen |
| 2B4 | BV650 | 2-69 | BD Pharmingen |
| CD2 | BV510 | 2H7 | BD Pharmingen |
| CD8 | BV570 | RPA-T8 | BIOLEGEND |
| CD57 | BB630 | NK-1 | BD Pharmingen |
| KLRG1 | PE-Dazzle594 | 2F1/KLRG1 | BIOLEGEND |
| PD1 | BV605 | EH12.1 | BD Pharmingen |
| TIM-3 | BV750 | 7D3 | BD Pharmingen |
| HLA-DR | BUV661 | G46-6 | BD Pharmingen |
| IL-7R | BUV805 | HIL-7R-M21 | BD Pharmingen |

**Supplemental Table 2**. 28-color panel for NK cell phenotyping
